# Vitronectin mediates survival of human WJ-MSCs under inflammatory temperature stress via cell cycle arrest

**DOI:** 10.1101/2022.02.23.481475

**Authors:** Umesh Goyal, Ashiq Khader C, Malancha Ta

## Abstract

Due to their anti-inflammatory and immunomodulatory capabilities, mesenchymal stem cells (MSCs) are being widely used in cell-based therapies for the treatment of a wide spectrum of inflammatory disorders. Despite their promises, substantial cell loss post transplantation leads to compromised therapeutic benefits in clinical trials, which remains a challenge to overcome. Inflammatory microenvironment comprises the presence of pro-inflammatory cytokines, elevated temperature, etc., which could hamper MSC viability following transplantation. Thus, identifying the underlying molecular factors controlling survival mechanism under such stress conditions, thereby, improving MSC survival becomes important for optimising MSC-based therapy. Also, since MSCs from different origins have significantly varied biology, choosing the appropriate MSC source could be crucial in determining the fate of transplanted MSCs in stressful milieu.

As extracellular matrix (ECM) components can mediate cell survival signals, in the present study, we have evaluated the role of ECM matricellular protein, vitronectin (VTN), in the survival of human umbilical cord-derived Wharton’s Jelly MSCs (WJ-MSCs) under the condition of inflammatory temperature stress. On exposure to 40°C, WJ-MSCs underwent cell cycle arrest with no significant change in viability status, along with an induction in VTN expression both at mRNA and protein levels. Interestingly, inhibition of pro-survival signalling pathways, ERK or PI3K, at 40°C led to further upregulation in VTN expression without any significant impact on viability or cell cycle arrest status. However, on knocking down VTN in WJ-MSCs at 40°C, decrease in viable population along with reversal of cell cycle arrest were noted. Moreover, inhibition of pro-survival pathways ERK or PI3K, in VTN knocked down WJ-MSCs at 40°C, led to a dramatic reduction in the viable population accompanied with reversal in cell cycle arrest.

Altogether, our findings highlighted the protective role of VTN in mediating survival of WJ-MSCs under inflammatory temperature stress condition, by possibly involving autophagy as an underlying mechanism.

## Introduction

Mesenchymal stem cells (MSCs) have emerged as a potential candidate for the treatment of inflammatory diseases like graft versus host disease (GVHD), Crohn’s disease, inflammatory bowel disease (IBD), etc. (1). This is due to their anti-inflammatory, immunomodulatory, and regenerative properties which are mediated via paracrine mechanism (1, 2). Hence, recently MSCs have been rechristened as ‘Medicinal Signaling Cells’ (3). Though many pre-clinical studies demonstrated overwhelmingly positive results for the use of MSCs for the treatment of various inflammatory associated diseases, however, when translated into human trials, similar scale of efficacy has been lacking (1). Thus, although MSC based therapy has been shown to be relatively safe, the efficacy of the treatment remains uncertain. The reduced efficacy is largely due to MSCs’ low survival post transplantation, arising as a result of the harsh microenvironment at the pathophysiological site (1). Thus, a better understanding of the interaction of MSCs with the pathological microenvironment is an essential step to fully unlock the potential of MSC based therapy. Additionally, the choice of the source of MSCs could be important in determining the clinical outcome of the treatment since MSCs obtained from different sources’ significantly differ in their biology (4). Bone marrow MSCs are the most frequently investigated source of MSCs in clinical trials. However, the umbilical cord can be a suitable and convenient alternative source of MSCs with several advantages, like superior proliferation ability and better immunomodulatory properties, over other adult tissue sources (5). Wharton’s jelly (WJ) is the mucoid, connective tissue surrounding the umbilical cord vessels and a rich source of MSCs (6).

Inflammation is a complex pathophysiological condition accompanied by prevalence of immune cells, pro-inflammatory cytokines, oxidative stress and elevated temperature (7). These micro-environmental factors at the inflammatory site can adversely affect the viability and functionality of MSCs. Hence, in order to improve and attain the maximum benefit out of MSC based therapy, it becomes crucial to scrutinize and evaluate the response of MSCs to inflammatory stress. Elevated temperature (fever) is one of the four cardinal signs and characteristics response of inflammation (8). However the effect of elevated temperature in the physiological fever range on the survival of MSCs and the survival strategies adopted has not been extensively studied.

Previous report from our group had demonstrated crosstalk between NF-κβ and p53 in G0/G1 arrested WJ-MSCs, when they were exposed to febrile temperature stress of 40°C (9). In yet another report from our lab, WJ-MSCs exhibited increased adhesion and reduced migration at 40°C with an underlying NF-κβ pathway and certain ECM genes and MMP1 being responsible (10).

Vitronectin (VTN) is a matricellular ECM glycoprotein present in large concentration in the serum and ECM, was first identified as an adhesive glycoprotein involved in the attachment and spreading of the cell (11). Subsequently, VTN expression has been shown to be induced in response to stress and inflammation, exhibiting tissue repair and remodelling activity (11–13). The importance of VTN in cell survival under various contexts also has been demonstrated earlier by a few reports. An earlier study had shown that VTN played a protective role towards TFG-β induced apoptosis of alveolar epithelial cells, via its interaction with integrins (14). The anti-apoptotic and proliferative role of VTN in nasopharyngeal carcinoma (NPC) in response to ionizing radiation by activation of ATM-Chk2 and ATR-Chk1 pathways has been shown (15). In human umbilical vein epithelial cells, VTN was reported to improve cell survival post-radiation injury by attenuating the expression of p21 and inhibiting apoptotic cell death (16). Yet another report had established the anti-apoptotic function of VTN in neutrophils via interaction of RGD domain to β1, β3, or β5 integrin (17). In the area of MSCs, a previous study from our group had shown VTN as an important candidate for maintaining WJ-MSCs’ viability under serum deprivation (18). However, as per our knowledge, not much has been reported on the importance of VTN in the survival of MSCs in the context of temperature stress.

Hence, in the present study, we have evaluated VTN’s role in the viability of WJ-MSCs under the condition of inflammatory temperature stress (40°C). We observed that WJ-MSCs exposed to 40°C stress exhibited an increase in VTN expression both at mRNA and protein levels, and underwent a G0/G1 cell cycle arrest with no significant impact on viability. Inhibition of pro-survival pathways, ERK or PI3K, at 40°C led to further upregulation in VTN expression with no significant change in cell viability and cell cycle status. Interestingly, esiRNA mediated knockdown of VTN in WJ-MSCs at 40°C with or without pro-survival pathway inhibitors resulted in reduction of cellular viability accompanied by reversal in G0/G1 cell cycle arrest. These results established that VTN acted as a protective factor and maintained the viability of WJ-MSCs at 40°C temperature stress by adopting G0/G1 cell cycle arrest mechanism. Finally, autophagy was indicated as a mechanism adopted by VTN in executing its pro-survival role.

## Materials and methods

Human umbilical cords were collected after full-term births (vaginal or caesarean delivery) with informed consent of the donor following the guidelines laid down by the Institutional Ethics Committee (IEC) and Institutional Committee for Stem Cell Research and Therapy (IC-SCRT) at IISER, Kolkata, India. WJ-MSCs were isolated from the perivascular region of the umbilical cord termed as Wharton’s jelly by explant culture method as described earlier (19). All experiments were performed between passages 4 to 6, with cell seeding density of 5000 cells/cm^2^. Cell were dissociated using TrypLE express (Life technologies), a gentle animal origin-free recombinant enzyme.

WJ-MSCs were initially plated in complete medium comprising of KnockOut DMEM (Dulbecco’s modified Eagle’s medium) supplemented with 10% fetal bovine serum (FBS), 2 mM Glutamine and 1X PenStrep (all from Life technologies) at 37°C for 24 h. Next, for temperature stress treatment, WJ-MSCs were exposed to 40°C for 48 h. For hypoxia treatment, cells were exposed to 2% oxygen for 48 h. For control culture conditions, cells were continued to grow at 37°C for another 48 h. Finally, at the end of 72 h, cells were harvested for respective experiments. For signalling pathway inhibition studies, cells were treated with pathway specific small molecule inhibitors and their respective vehicle controls under 40°C or hypoxia condition for a period of 48 h. PI3K pathway inhibitor, LY294002, ERK pathway inhibitor, FR180204 and NF-κβ pathway inhibitor, BAY 11-7082 were used at an optimized concentrations of 30 μM, 30 μM and 4 μM respectively (all from Sigma-Aldrich). For autophagy inhibition studies, chloroquine (Sigma-Aldrich) was used at a concentration of 25 μM, as an autophagy specific inhibitor.

### siRNA transfection

For siRNA transfection WJ-MSCs were plated and grown under control conditions till 50-60% of confluency was achieved. Next, transfection was performed using Lipofectamine 3000 in Opti-MEM I (both from Life technologies) medium as per manufacturer’s protocol. Endonuclease-treated siRNA pool (esiRNA) generated against *VTN* was used for VTN knockdown, while for experimental control MISSION^®^ siRNA Universal Negative Control (both from Sigma Aldrich) was used at a concentration of 30 nm each. Following 12-14 h of transfection WJ-MSCs were exposed to 40°C in the absence or presence of pathway specific small molecule inhibitors for 36-48 h. Next cells were harvested and analysis was performed.

### Population doubling and doubling time

The number of population doublings was calculated using the formula: PD= [log_10_ (NH) – log_10_ (NI)]/log_10_ (2), while the population doubling time was obtained by the formula: PDT =t * log_10_ 2 / (log_10_NH - log_10_NI). NI: the inoculum cell number; NH is the cell harvest number, and t is the time of the culture (in h).

### MTT assay

To assess cell proliferation, 3-(4, 5-dimethylthiazol-2-yl)-2, 5-diphenyltetrazolium bromide (MTT) assay was performed using standard protocol. In brief, WJ-MSCs were seeded onto a 96-well plate and exposed to different treatments. Next MTT (Sigma Aldrich) dye was added to each well at a final concentration of 0.5 mg/mL and incubated at 37°C. The reaction was terminated by adding dimethyl sulfoxide (DMSO; Merck) to solubilize purple-colored formazan crystals. Absorbance was measured at 595 nm using a plate reader (Epoch). Percent change in proliferation was calculated with respect to control after subtracting the background absorbance. Each assay was carried out in triplicate.

### Cell cycle analysis

In order to assess the change in the cell cycle phase distribution following different treatments, control and treated cells were harvested and fixed in 70 % ethanol in PBS at - 20°C overnight. The following day, cells were incubated with 50 μg/ml propodium iodide (PI) (Sigma Aldrich) containing 100 μg/ml RNAse A (Thermo Fischer Scientific) at 37°C. Samples were analysed by BD LSRFortessa™ Cell Analyzer flow cytometer, and the fraction of cells in the different phases was assessed using BD FACSDiva™ Software (BD Biosciences).

### Apoptosis assay: Annexin V-PI staining

To assess the viability of WJ-MSCs following different treatments, floating and adherent populations of cells were harvested, and apoptosis assay was performed using Alexa Fluor 488 Annexin-V/Dead Cell Apoptosis Kit (Life Technologies) as per the manufacturer’s instructions. Percentage of the viable and apoptotic cells were analysed by flow cytometry (BD LSRFortessa™ Cell Analyzer; BD Biosciences). Ten thousand events were analysed in each of the samples. Compensation controls were included in every experiment.

### Western blotting

To compare the relative expression of protein following different treatments, WJ-MSCs were lysed in RIPA buffer containing protease inhibitor cocktail (Sigma Aldrich) and phosphatase inhibitor (Abcam). Protein concentration was quantified using the Bradford reagent (Bio-Rad). SDS-PAGE gel electrophoresis and Western blotting were performed as per standard protocols. The primary and secondary antibodies used were anti-VTN, anti-p53, anti-GAPDH (all from Santa Cruz Biotechnology, Inc.), anti-p62 (GeneTex) and horseradish peroxidase (HRP)-linked anti-mouse IgG (Cell Signaling Technology, Inc.) respectively.

### Immunofluorescence

Immunostaining of control and treated WJ-MSCs was performed as per standard protocol. Briefly, cells grown on coverslips were fixed in 4% paraformaldehyde, followed by blocking and permeabilization with 0.1% Triton X-100. Next, the primary antibody incubation was done overnight at 4°C, following which the secondary antibody incubation was carried at room temperature for 1 h. After labeling the nucleus with DAPI (Sigma Aldrich), coverslips were mounted with VECTASHIELD antifade mounting medium (Vector Laboratories). The primary antibodies used were anti-VTN (Santa Cruz Biotechnology, Inc.), and anti-vimentin (VIM) (Cell Signalling Technology). The secondary antibodies used were goat anti-mouse IgG H&L (Alexa Fluor^®^ 488) preadsorbed (Abcam) and goat anti-rabbit IgG (H+L) cross-adsorbed secondary antibodies, Alexa Fluor 568 (Thermo Fischer Scientific), respectively. Images were acquired at 40X magnification using Zeiss Apotome module microscope using Zen software. For quantification of VTN expression under different treatments, region of interest (ROI) was manually demarcated and mean fluorescence intensity of the cell was quantified using Image J (NIH) software. Background mean fluorescence intensity was subtracted to obtain the corrected mean fluorescence intensity.

### RNA isolation and cDNA synthesis

WJ-MSCs were lysed in Tri-Reagent (Sigma-Aldrich), and total RNA was isolated as per the manufacturer’s protocol. Next, RNA yield was quantified using Nanodrop 2000 spectrophotometer (Thermo Fischer Scientific). Equal amount of RNA was taken and cDNA synthesis was performed using Verso cDNA synthesis kit (Thermo Fischer Scientific) according to the manufacturer’s instructions.

### Quantitative reverse transcription-polymerase chain reaction (qRT-PCR)

The changes in gene expression following different treatments were quantified via qRT-PCR using specific primers. All the reactions were performed with PowerUp SYBR™ Green Master mix (Applied Biosystems) using the CFX96 Touch Real-Time PCR System (Bio-Rad). Fold change in expression was calculated using the 2^-ΔΔCT^ method. Melting curve analysis was done to confirm single, specific PCR products.

GAPDH was considered as the endogenous control and used to normalize gene expression levels. The gene accession number, primer sequences and amplicon sizes are listed in **Table 1**.

**Table 1:**
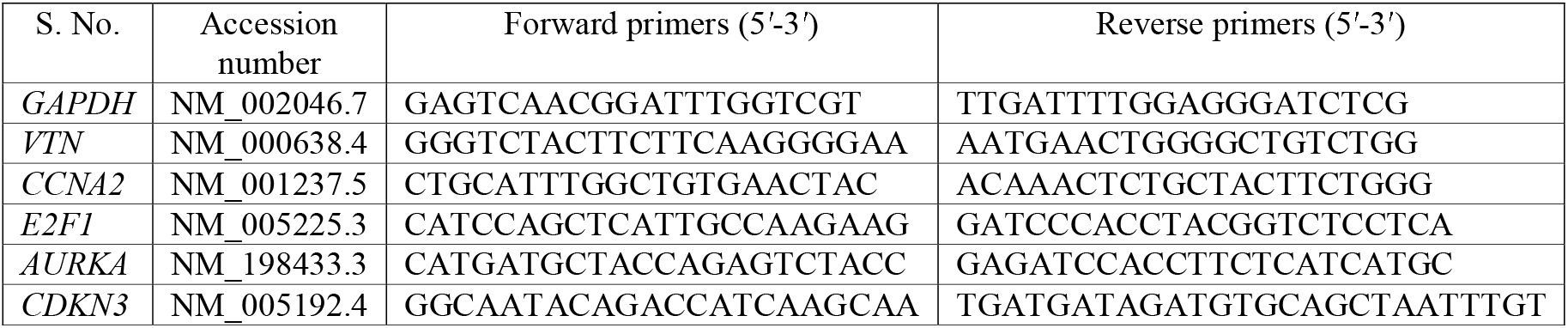
Primer sequences used for quantitative RT-PCR.

### Statistical analysis

Data analysis and graphical representations were performed using GraphPad Prism 5/8 software (GraphPad). All data are presented as mean ± SEM. The analytical methods used were the Student’s two-tailed t test, one-way ANOVA, and two-way ANOVA. In multigroup analysis, ANOVA was followed by Tukey’s or Bonferroni’s test. Significance was confirmed at *p* < 0.05, 0.01 or 0.001 levels. Each experiment was performed with at least 3 independent biological samples.

## Results

### Impact of 40°C on VTN expression in WJ-MSCs

WJ-MSCs exposed to 40°C stress showed an increase in VTN expression both at mRNA (*p* < 0.01) and protein levels (not significant) (**Fig. 1A**, **B**). Immunofluorescence staining performed to examine the subcellular distribution of VTN at 40°C depicted an increased expression of VTN in the nucleus and cytoplasm compared to control WJ-MSCs (**Fig. 1C**). Additionally, we also saw an elevated level of VTN in the ECM of 40°C treated WJ-MSCs. Co-staining with mesenchymal marker vimentin showed positive reactivity confirming the mesenchymal nature of WJ-MSCs under different treatments (**Fig. 1C**). Further, on examining the temporal expression pattern of VTN under 40°C keeping the cell seeding density constant, WJ-MSCs were shown to exhibit an increase in VTN protein expression in a time-dependent manner under 40°C (**Fig. 1D**). Similarly, WJ-MSCs plated at increasing cell seeding densities of 2000, 5000, and 8000 cells/cm^2^, and exposed to 40°C for 48 h, demonstrated an increase in VTN protein expression in a confluency dependent manner (**Fig. 1E**).

**Figure 1:**
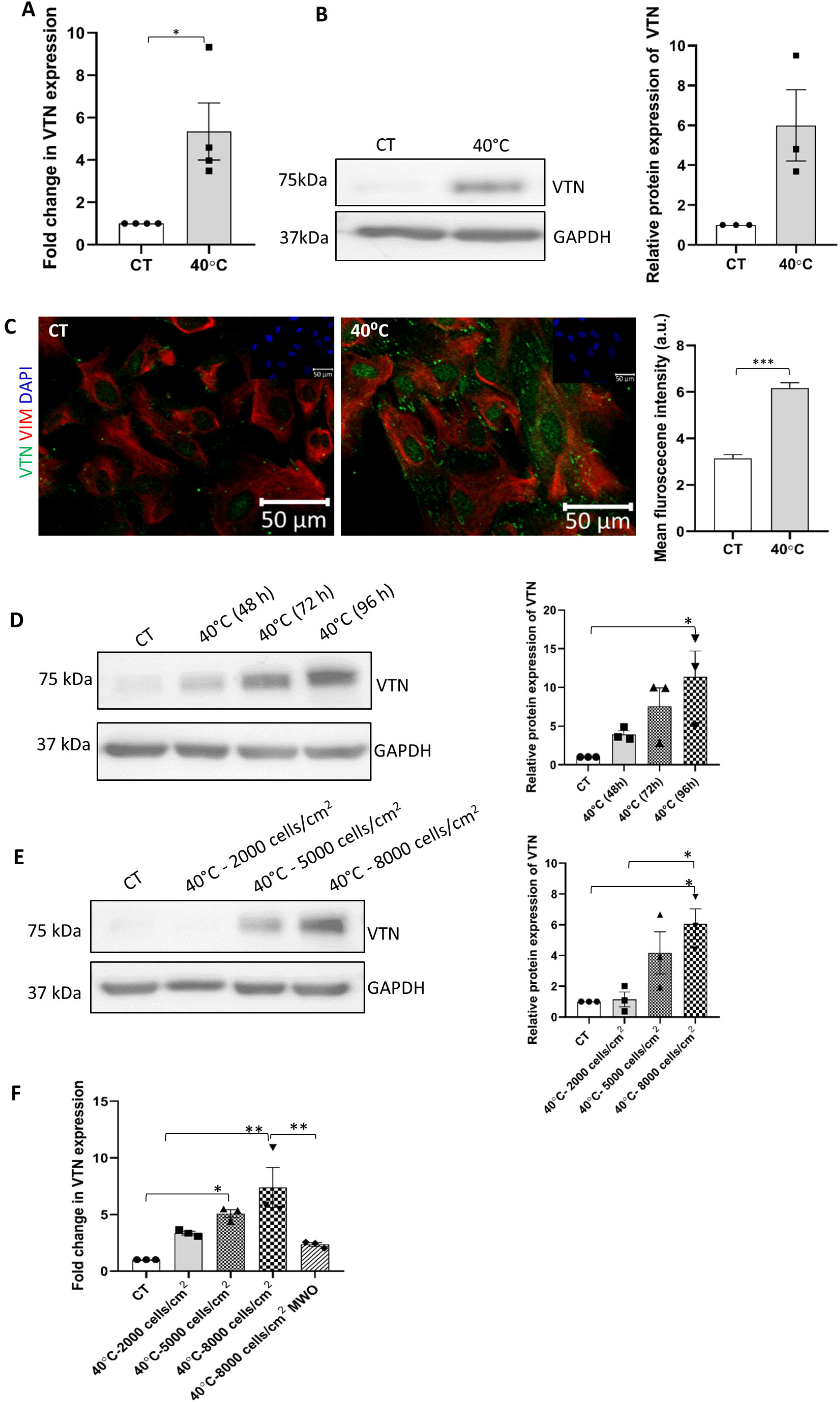
Impact of 40°C temperature stress on VTN expression in WJ-MSCs. (**A**) mRNA expression of *VTN* as detected by qRT-PCR. *GAPDH* was used as an endogenous control to normalise the gene expression. (**B**) Representative Western blot image depicting VTN protein expression. Band density of protein expression levels of VTN was quantified relative to GAPDH, which was used as loading control, and plotted. (**C**) Representative immunofluorescence images depict distribution of VTN (green) and VIM (red) in control and 40°C treated WJ-MSCs. Nucleus was labelled with DAPI (blue) shown in inset. Mean fluorescence intensity for VTN expression was quantified per cell using Image J software and plotted (n = 80) from three different biological samples. Change in VTN protein expression in exposure time and seeding density dependent manner. (**D**) WJ-MSCs were exposed to 40°C for durations of 48, 72, and 96 hs and VTN protein expression was evaluated. Representative Western blot image depicting VTN protein expression is displayed. Band density of VTN was quantified and plotted relative to GAPDH, used as loading control. (**E**) WJ-MSCs were plated at different cell seeding density of 2000, 5000 and 8000 cells/cm^2^ and exposed to 40°C for a period of 48 h. VTN protein expression was evaluated by Western blotting, representative image is shown from three independent biological samples. Band density of VTN was quantified and plotted relative to GAPDH, used as loading control. (**F**) Effect of medium washout (MWO) on *VTN* mRNA expression. WJ-MSCs plated at high seeding density of 8000 cells/cm^2^ under 40°C, when subjected to medium change every 12 h showed reduction in the *VTN* mRNA expression as compared to the MSC samples without any medium change. *GAPDH* was used as an endogenous control to normalise the gene expression. Each bar represents mean ± SEM. * represent *p* < 0.05, ** represents *p* ≤ 0.01, and *** represents *p* ≤ 0.001. Data shown are representative of at least three independent biological samples (n ≥ 3).

Induction in VTN protein expression with increasing seeding density might be attributed to an altered gene expression program of VTN, due to autocrine/paracrine feedback signalling via secreted VTN or other soluble factor/s in the conditioned medium. In order to confirm the hypothesis we performed a medium washout (MWO) experiment. WJ-MSCs plated at the highest cell seeding density of 8000 cells/cm^2^ were exposed to 40°C for a period of 48 h and subjected to medium change every 12 h. As hypothesized, the cells subjected to the above MWO treatment, exhibited a strong downregulation in *VTN* mRNA expression (*p* < 0.01) (**Fig. 1F**).

### Effect of pro-survival pathways’ inhibition on VTN expression, viability and cell cycle status of WJ-MSCs under 40°C

To explore the underlying molecular mechanism for regulation of VTN expression under 40°C condition, we checked for VTN expression at mRNA and protein levels on treating WJ-MSCs with specific small molecule inhibitors of different signalling pathways under 40°C. Inhibition of ERK and PI3K pathways exhibited a further upregulation in *VTN* mRNA expression, while a reduction in the expression was noted with NF-κβ pathway inhibition (not significant) (**Fig. 2A**). Similarly, a strong induction in VTN protein expression was observed with the inhibition of ERK or PI3K signalling pathways under 40°C, though not significant, as compared to only 40°C treated WJ-MSCs (**Fig. 2B**).

**Figure 2:**
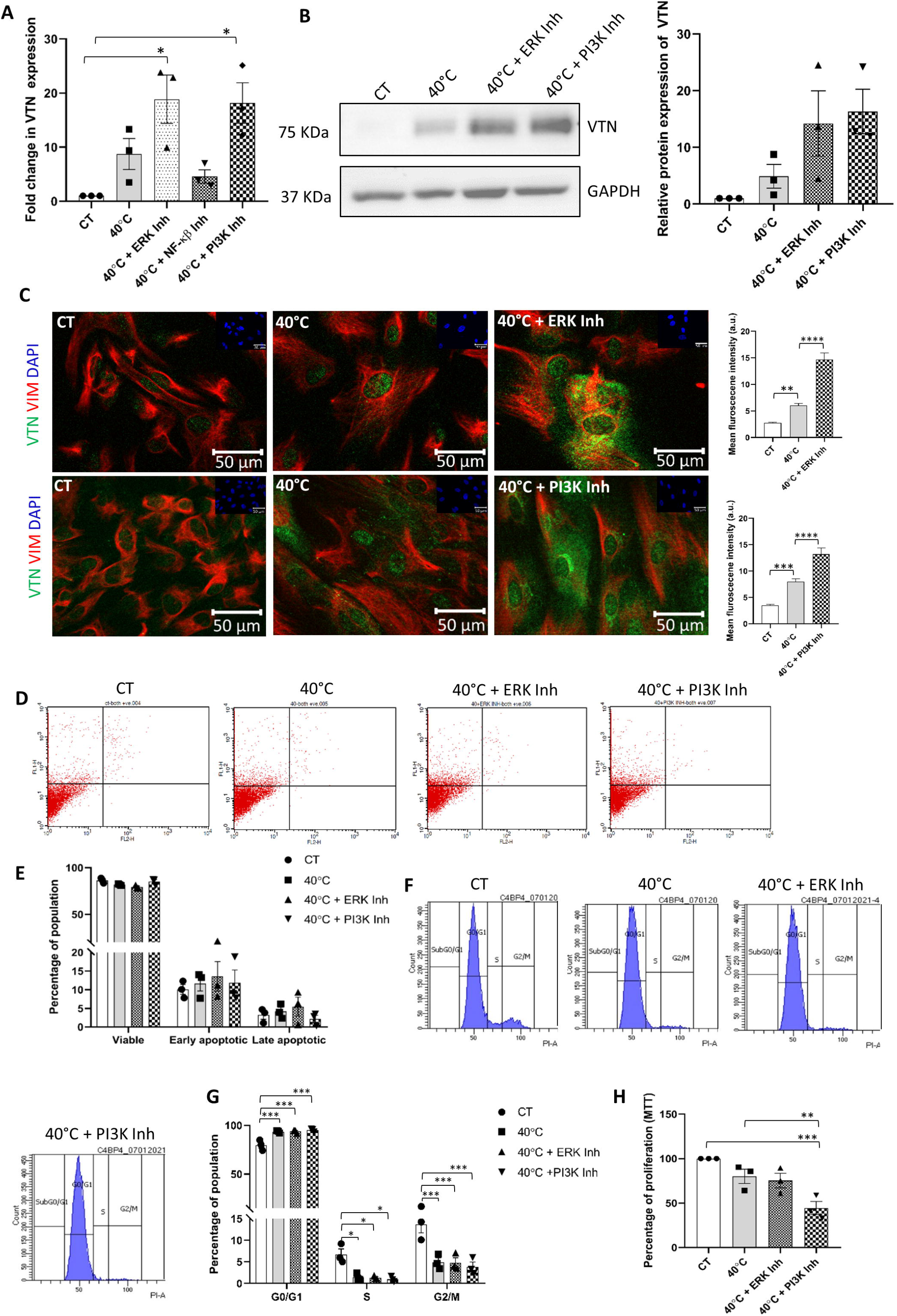
Effect of inhibition of signalling pathways on VTN expression, cell cycle profile and viability status of WJ-MSCs under the 40°C temperature stress. WJ-MSCs were exposed to 40°C in the absence or presence of small molecule inhibitors for certain signalling pathways for 48 h. (**A**) *VTN* mRNA expression was detected by qRT-PCR. *GAPDH* was used as an endogenous control to normalise the gene expression. (**B**) VTN protein expression was detected in WJ-MSCs treated at 40°C for 48 h in the absence or presence of ERK pathway inhibitor FR180204 or PI3K pathway inhibitor, LY294002. Representative Western blotting images from three independent biological samples are displayed. Band densities were quantified and plotted relative to GAPDH expression used as loading control. (**C**) Representative immunofluorescence images depict the distribution of VTN (green) and VIM (red) in WJ-MSCs treated at 40°C for 48 h in the presence or absence of ERK or PI3K pathway inhibitor, and under control conditions. Nuclei stained with DAPI (blue) are shown in inset. Mean fluorescence intensity for VTN expression per cell was quantified and plotted for a total of 45 cells for ERK pathway inhibition from two different biological samples, and 65 cells for PI3K pathway inhibition from three different biological samples. (**D**) Detection of viable population via annexin-V-PI staining of WJ-MSCs treated at 40°C in the absence or presence of ERK or PI3K pathway specific small molecule inhibitor and under control condition. (**E**) Histogram represents the percentages of viable, early apoptotic, and late apoptotic populations in control and treated WJ-MSC samples. (**F**) Flow cytometry-based cell cycle analysis of WJ-MSCs exposed to 40°C in the presence or absence of ERK or PI3K pathway specific inhibitor and under control conditions are demonstrated. Data shows the distribution of WJ-MSCs in the G0/G1, S and G2M phases. (**G**) Percentages of cells in each phase of the cell cycle for the indicated treatment conditions are compared and represented by a histogram. (**H**) MTT assay comparing proliferation at 40°C, for the above mentioned treatments and control condition are shown. Each bar represents mean ± standard error mean. * represent *p* ≤ 0.05, ** represent *p* ≤ 0.01, *** represents *p* ≤ 0.001 and **** represents *p* ≤ 0.0001. Data shown are representative of at least three independent biological samples (n ≥ 3).

As an increase in the expression of VTN, both at mRNA and protein levels, was noted with inhibition of ERK and PI3K signalling pathways individually, we next performed immunofluorescence staining to confirm the increase in VTN expression and its sub-cellular distribution. Immunofluorescence staining data revealed a widespread increase in VTN expression with ERK or PI3K pathway inhibition at 40°C, as compared to untreated 40°C samples. This increase was observed in the nucleus, cytoplasm, as well as the ECM (**Fig. 2C**). Co-staining with mesenchymal marker vimentin showed positive reactivity confirming the mesenchymal nature of WJ-MSCs under different treatments (**Fig. 2C**).

Effect on cell viability was evaluated next using annexin-V-PI staining followed by flow cytometry. WJ-MSCs exposed to 40°C stress did not show any apoptotic change and cell viability was not found to be significantly impacted (**Fig. 2D, E**). Furthermore, inhibition of pro-survival signalling pathways, ERK and PI3K, individually, at 40°C also did not demonstrate any significant change in the apoptotic population or the viability status of WJ-MSCs as compared to only 40°C treated MSCs. (**Fig. 2D, E**).

However, exposing WJ-MSCs to 40°C stress led to increased accumulation of cells in the G0/G1 phase from 79.69 ± 3.07 to 93.64 ± 0.70 (*p* < 0.001) with a corresponding decline in the S and G2/M phase populations from 6.7 ± 1.26 to 1.44 ± 0.48 (*p* < 0.05) and 13.69 ± 1.94 to 4.9 ± 0.99 (*p* < 0.001), respectively, as compared to control WJ-MSCs (**Fig. 2F, G**). Further, on inhibiting ERK and PI3K pathways individually, using specific small-molecule inhibitors, FR180204 and LY294002, respectively, at 40°C, WJ-MSCs continued to be in G0/G1 arrested phase with a concomitant reduction in the S and G2/M phase and no further change in the cell cycle profile was observed (**Fig. 2F, G**). Moreover, assessment of proliferation using MTT assay showed a reduction in proliferation of WJ-MSCs at 40°C, though not significant. Treatment with ERK pathway inhibitor under 40°C didn’t show any further change, while, PI3K pathway inhibition under 40°C led to a significant reduction (*p* < 0.01) in the proliferation as compared to only 40°C exposed WJ-MSCs (**Fig. 2H**).

### VTN knockdown led to reduction in cell viability with reversal in G0/G1 cell cycle arrest

Apoptosis analysis in the previous set of experiments suggested that the induced VTN expression might have provided a protective advantage to WJ-MSCs exposed to 40°C stress condition. Hence, in order to confirm this, VTN was knocked down with *VTN* specific esiRNA in WJ-MSCs followed by exposure to 40°C for 48 h, and viability and cell cycle analysis were assessed. VTN knocked down WJ-MSCs demonstrated downregulation in *VTN* mRNA expression (*p* < 0.01) (**Fig. 3A**). This resulted in a reduction in viable population of WJ-MSCs from 80.20 ± 3.27 to 69.82 ± 3.85, though not significant, as compared to NC siRNA transfected WJ-MSCs. Also, a corresponding increase in early and late apoptotic populations from 13.83 ± 3.03 to 17.97 ± 6.5 and 5.26 ± 2.46 to 11.29 ± 3.73, respectively, again not significant, was noted from apoptosis assays (**Fig. 3B, C**).

**Figure 3:**
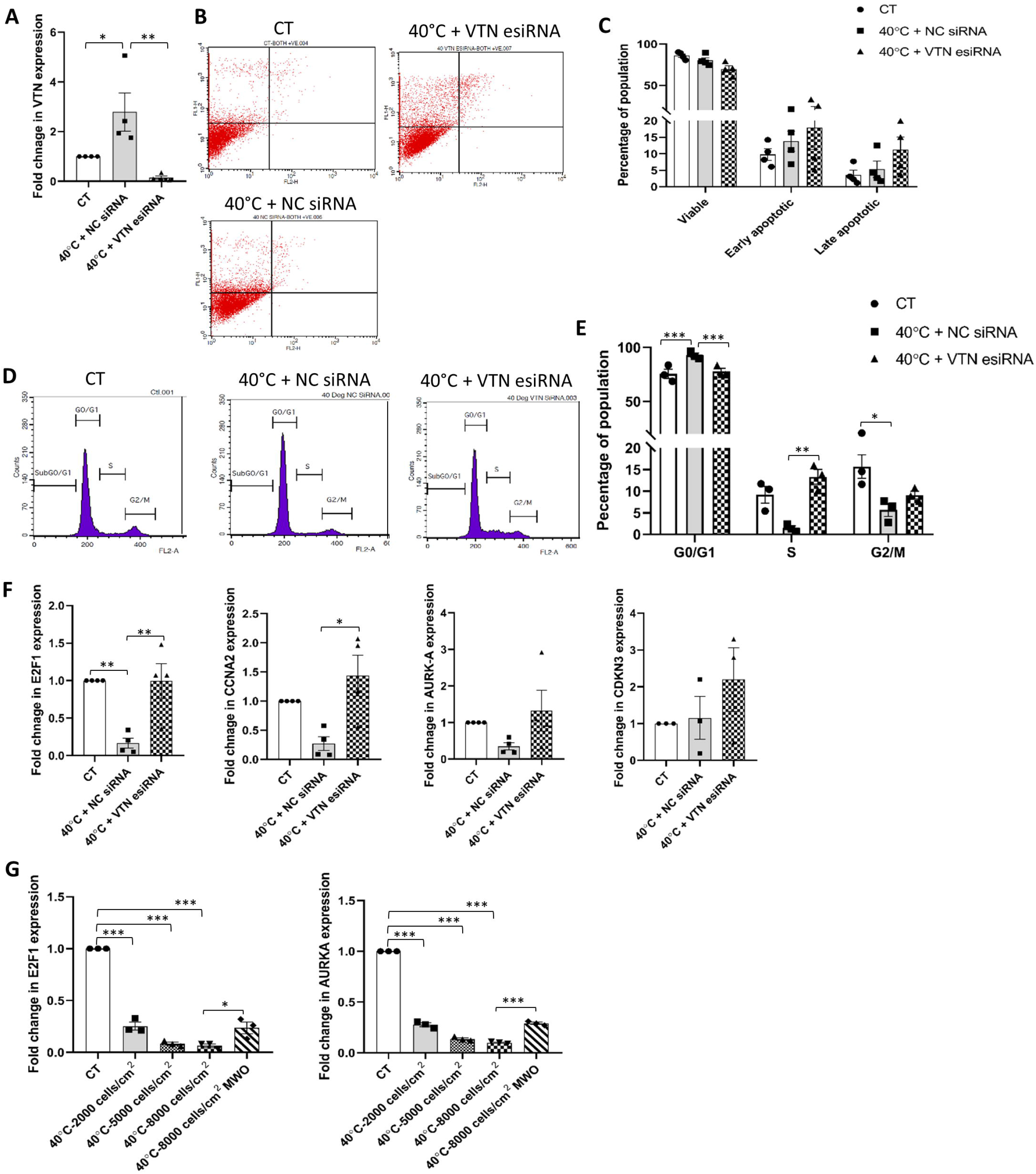
Impact of VTN knockdown on viability and cell cycle status of WJ-MSCs exposed to 40°C temperature stress. WJ-MSCs transfected with *VTN* esiRNA, or NC siRNA were exposed to 40°C for 48 h. (**A**) qRT-PCR analysis of *VTN* mRNA expression was performed, with *GAPDH* as an endogenous control to normalise the gene expression. (**B**) Assessment of viability status by annexin-V-PI staining followed by flow cytometry analysis. Representative flow cytometry data from at least three different biological samples are displayed. (**C**) Quantitation and comparison of percentages of viable, early apoptotic and late apoptotic populations between NC siRNA or *VTN* esiRNA transfected WJ-MSCs at 40°C and control WJ-MSCs cultures are depicted by the histograms. (**D**) Cell cycle analysis of WJ-MSCs treated with NC siRNA or *VTN* esiRNA at 40°C and control WJ-MSCs. (**E**) Percentages of cells in different phases of the cell cycle are shown by the histogram. (**F**) mRNA expression levels of cell cycle markers *E2F1, CCNA2, AURKA* and *CDKN3* in NC siRNA or *VTN* esiRNA transfected WJ-MSCs under 40°C, are demonstrated by qRT-PCR analysis. *GAPDH* was used as an endogenous control to normalise the gene expression. (**G**) Effect of medium washout (MWO) on the expression of cell cycle markers, *E2F1* and *AURKA. GAPDH* mRNA expression was considered as an endogenous control. Each bar represents mean ± SEM. * represent *p* ≤ 0.05, ** represents *p* ≤ 0.01 and *** represents *p* ≤ 0.001. Data shown are representative of at least three independent biological samples (n ≥ 3).

Further to confirm whether the reduction in percentage of viable WJ-MSCs was accompanied with the reversal in cell cycle arrest seen under 40°C, cell cycle profile was evaluated. *VTN* esiRNA transfection resulted in a rescue in G0/G1 arrested cell population, bringing down the percentage from 92.85 ± 1.99 to 77.82 ± 2.77 (*p* < 0.001), compared to NC siRNA transfected WJ-MSCs under 40°C (**Fig. 3D, E**). There was also a corresponding increase in the S phase cells from 1.46± 0.48 to 13.32 ± 1.68 (*p* < 0.01), and G2/M phase cells from 5.66 ± 1.48 to 9.09 ± 0.89 (not significant) in VTN knocked down WJ-MSCs (**Fig. 3D, E**). Next, to validate the reversal in G0/G1 arrest, mRNA expression levels of cell cycle markers involved in different phases of the cell cycle were investigated. The reduction observed in the expression of cell cycle progression markers *E2F1, CCNA2, AURKA*, and *CDKN3* at 40°C was found to be rescued in VTN knocked down WJ-MSCs at 40°C (**Fig. 3F**). This further confirmed the G0/G1 cell cycle phase arrest reversal with knockdown of VTN under 40°C.

Since we had observed a reduction in *VTN* expression in the cell density MWO experiment, we next also evaluated the mRNA expression level of cell cycle progression markers like *E2F1* and *AURKA* in these samples. It was seen that as *VTN* mRNA expression increased in the MSCs with increasing cell density at 40°C, *E2F1* and *AURKA* showed a marked reduction in their expression levels (*p* < 0.001) (**Fig. 3G**). However, this reduced expression of the cell cycle markers was rescued partially following washout of medium (*p* < 0.05) (**Fig. 3G**). Along with the rescue in the expression of cell cycle markers, there was an increase in population doublings and a concomitant decrease in population doubling time (**Supplementary fig. 1A, B**). This further demonstrated an inverse correlation between the expression of VTN and cell cycle progression status.

### Pro-survival pathway inhibition in VTN knocked down WJ-MSCs at 40°C led to significant level of apoptosis with reversal in G0/G1 arrest

Earlier, we had seen that there was no additional change in apoptosis level in WJ-MSCs, even on inhibiting survival pathways like ERK and PI3K under 40°C. This could be attributed to the increased expression levels of VTN in cells under ERK and PI3K pathway inhibition at 40°C. Therefore, to affirm the plausible role of VTN in maintaining the viability and cell cycle profile of WJ-MSCs under 40°C with inhibition of pro-survival PI3K pathway, WJ-MSCs were transfected with either *VTN* esiRNA or NC siRNA, and then exposed to 40°C for period of 48 h in presence or absence of PI3K pathway specific small molecule inhibitor, LY294002. *VTN* esiRNA treated WJ-MSCs showed a reduction in *VTN* mRNA expression (*p* < 0.01) when assayed for knockdown confirmation (**Fig. 4A**). Further, on assessing cell viability, *VTN* esiRNA transfected WJ-MSCs treated with LY294002, at 40°C, showed a significant decrease in the percentage of viable population from 88.5 ± 2.66 to 73.80 ± 3.20 (*p* < 0.001), with an increase in early and late apoptotic populations from 7.03 ± 2.09 to 13.30 ± 4.00 (not significant), and 2.93 ± 1.0 to 12.16 ± 2.3 (*p* < 0.05), respectively, as compared to NC siRNA transfected WJ-MSCs treated with LY294002, at 40°C (**Fig. 4B, C)**. Moreover, this decrease in viability was accompanied by a reversal in the G0/G1 cell cycle phase arrest. *VTN* esiRNA transfected WJ-MSCs treated with LY294002, at 40°C showed a reduction in the percentage of G0/G1 arrested cells from 94.5 ± 1.00 to 77.06 ± 8.15 (*p* < 0.05) with a concomitant increase in the S and G2/M phase cells from 1.83 ± 0.6 to 15.03 ± 8.28, and 3.6 ± 0.40 to 7.6 ± 0.635, respectively, though not significant, as compared to NC siRNA transfected WJ-MSCs under the same treatment (**Fig. 4D, E**).

**Figure 4:**
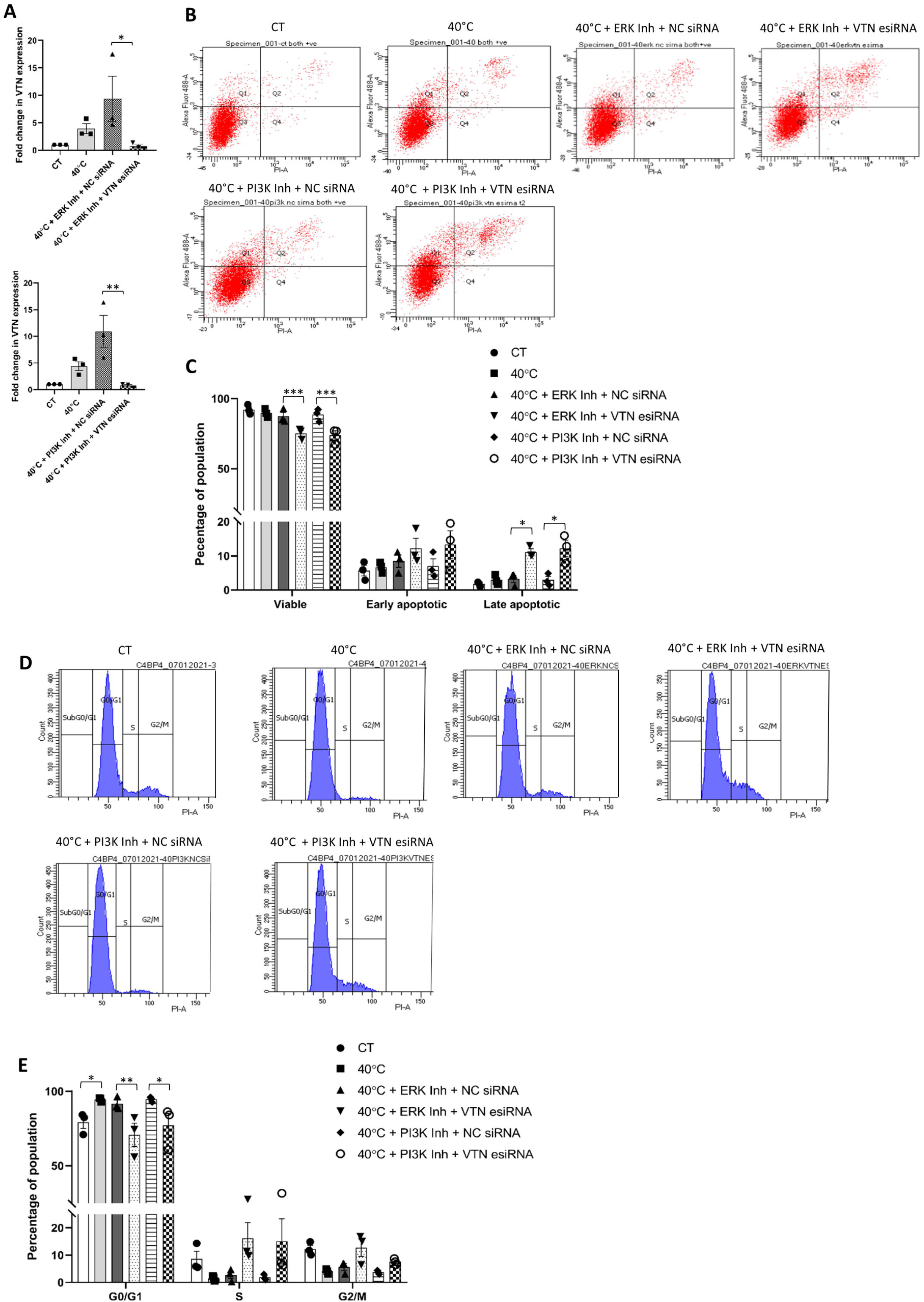
Influence of ERK or PI3K pathway inhibition on VTN transfected WJ-MSCs under 40°C temperature stress. WJ-MSCs transfected with *VTN* esiRNA, or NC siRNA were treated with ERK pathway inhibitor, FR180204 or PI3K pathway inhibitor, LY294002, at 40°C for 48 h. (**A**) *VTN* knockdown was validated at mRNA level as detected by qRT-PCR. *GAPDH* was used as an endogenous control to normalise the gene expression. (**B**) Assessment of viability status by annexin-V-PI staining followed by flow cytometry analysis. Representative flow cytometry data for apoptosis analysis from at least three independent biological samples are exhibited. (**C**) Comparison of percentages of viable, early apoptotic, and late apoptotic populations between control and indicated treatment conditions are depicted by the histogram. (**D**) Treated cells were stained with propidium iodide, and DNA content was analysed by flow cytometry for cell cycle analysis. Representative cell cycle analysis data using flow cytometry are shown for the indicated treatment conditions. (**E**) Percentages of cells in each phase of the cell cycle under control and indicated treatment conditions are depicted by a histogram.Each bar represents mean ± SEM. * represent *p* ≤ 0.05, ** represent *p* ≤ 0.01, and *** represents *p* ≤ 0.001. Data shown are representative of at least three independent biological samples (n > 3).

Similarly, WJ-MSCs transfected with *VTN* esiRNA or NC siRNA, were exposed to 40°C for a period of 48 h in presence or absence of ERK pathway specific small molecule inhibitor FR180204. *VTN* esiRNA transfected WJ-MSCs showed reduction in *VTN* mRNA expression (*p* < 0.05) when compared to NC siRNA transfected WJ-MSCs, as knockdown validation (**Fig. 4A**). Further, on assessing viability, *VTN* esiRNA transfected WJ-MSCs treated with FR180204, at 40°C, showed a significant decrease in the percentage of viable population from 87.36 ± 2.83 to 75.10 ± 2.77 (*p* < 0.001) as depicted by flow cytometry analysis. A corresponding increase in percentages of early and late apoptotic population from 8.43 ± 1.802 to 12.23 ± 2.91 (not significant) and 3.26 ± 1.084 to 11.10 ± 0.95 (*p* < 0.05) respectively, was also noted as compared to NC siRNA transfected WJ-MSCs under the same treatment (**Fig. 4B, C)**. Furthermore, decrease in viability was accompanied with a reversal in the G0/G1 cell cycle phase arrest. *VTN* esiRNA transfected WJ-MSCs treated with FR180204 at 40°C, showed reduction in the enrichment of G0/G1 cells from 91.60 ± 2.47 to 70.73 ± 7.85 (*p* < 0.01), with a concomitant increase in the percentages of S and G2/M phase cells from 2.73 ± 1.29 to 16.20 ± 5.66, and 5.6 ± 1.25 to 12.73 ± 3.2, respectively, though not significant, as compared to NC siRNA transfected WJ-MSCs under the same treatment (**Fig. 4D, E**).

### VTN expression, its regulation and role in viability under hypoxia stress condition

Though oxygen tension of 21% is used as standard cell culture practice, the physiological oxygen tension in stem cell niche is much lower, and as a result, MSCs are known to thrive well under hypoxia microenvironment which supports their stemness and proliferation (20). Thus, to understand the role of VTN, if any, in the survival of WJ-MSCs under hypoxic microenvironment, we aimed to study its expression pattern and regulation. Hypoxia treated WJ-MSCs displayed morphology similar to control WJ-MSCs (**Fig. 5A**). Further, growth kinetics assessment demonstrated no significant change as compared to control WJ-MSCs. The number of population doublings of 2.742 ± 0.143 and 2.802 ± 0.16 (**Fig. 5B**) and population doubling time of 26.39 ± 1.36 and 25.88 ± 1.53 h (**Fig. 5C**) were noted for control and hypoxia treated WJ-MSCs, respectively. Corresponding to these, VTN expression level of WJ-MSCs under hypoxia condition was also similar to control condition (**Fig. 5D, E**).

**Figure 5:**
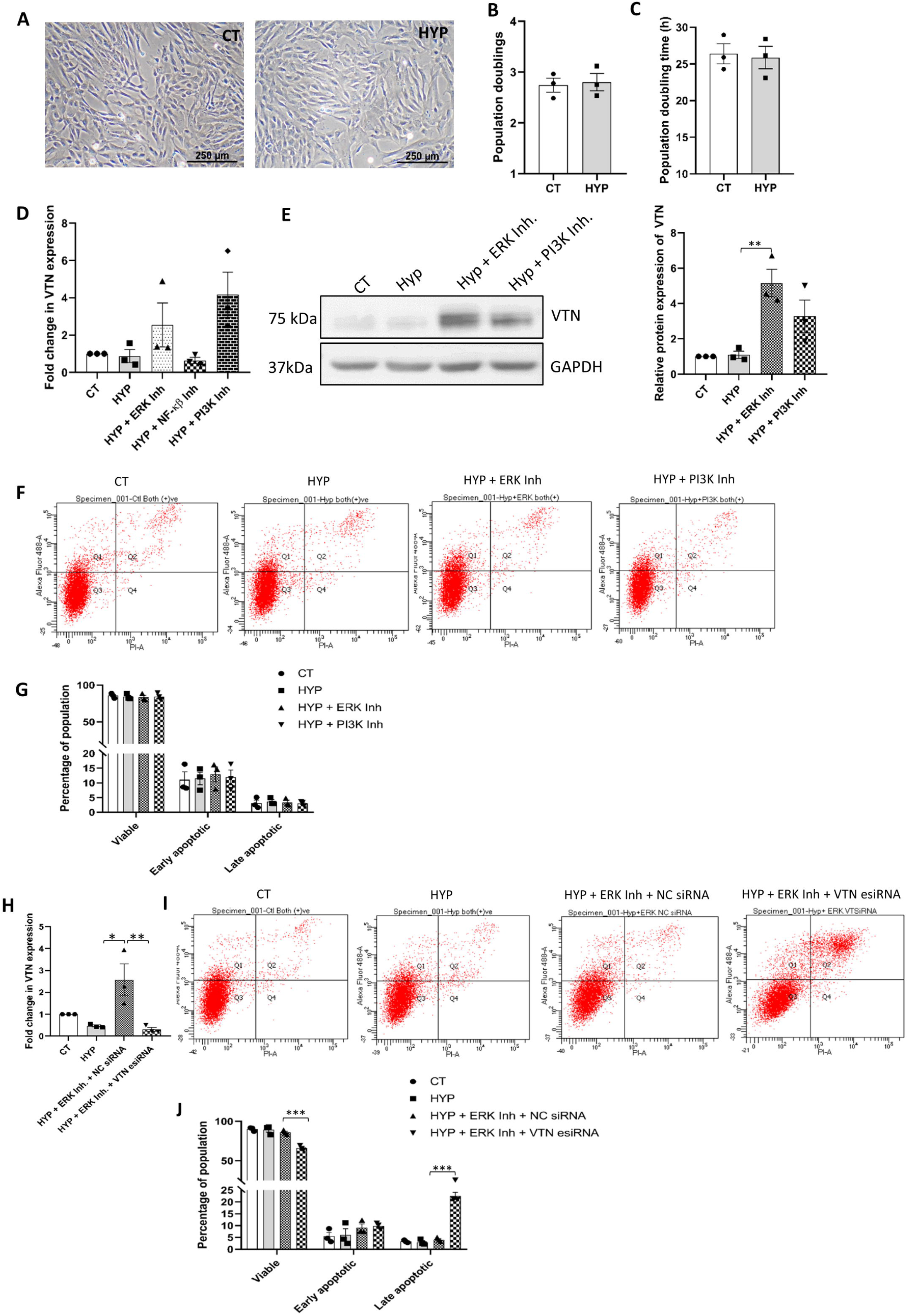
VTN expression, regulation and its role in viability under hypoxia condition. WJ-MSCs were exposed to hypoxia (2% O_2_) stress for 48 h. (**A**) Representative phase-contrast morphology images of the control and hypoxia treated WJ-MSCs are presented from three independent biological samples (10X magnification). Comparison of (**B**) number of population doublings and (**C**) population doubling time between control and hypoxia treated WJ-MSCs. (**D**) WJ-MSCs were exposed to hypoxia condition for 48 h in the absence or presence of indicated signalling pathway specific small molecule inhibitors. Next, the mRNA expression of *VTN* was evaluated by qRT-PCR. *GAPDH* was used as an endogenous control to normalise the gene expression. (**E**) VTN protein expression under hypoxia stress in absence or presence of ERK or PI3K pathway specific small molecule inhibitor was represented by Western blotting image from three independent biological samples, as shown. Band densities were quantified relative to GAPDH protein expression, which was used as loading control and plotted. (**F**) Detection of viability via flow cytometry analysis of WJ-MSCs exposed to hypoxia stress in the presence or absence of ERK or PI3K pathway inhibitors FR180204 and LY294002, respectively. Representative flow cytometry data for apoptosis analysis are shown from three independent biological samples. (**G**) Comparison of the viable, early, and late apoptotic populations under the condition of hypoxia in the presence or absence of ERK and PI3K pathway inhibitors and control condition are demonstrated by histogram. WJ-MSCs transfected with *VTN* esiRNA or NC siRNA were treated with ERK pathway inhibitor FR180204 under hypoxia stress for 48 h. (**H**) Reduction in *VTN* mRNA expression was confirmed by qRT-PCR. *GAPDH* was used as an endogenous control to normalise the gene expression. (**I**) Detection of viability by annexin-V-PI staining followed by flow cytometry analysis. Representative flow cytometry data are shown for apoptosis analysis from three independent biological samples. (**J**) Quantification and comparison of viable, early and late apoptotic populations between NC siRNA or *VTN* esiRNA transfected WJ-MSCs treated with ERK pathway inhibitor under hypoxia condition and control WJ-MSCs are demonstrated by a histogram. Each bar represents mean ± SEM. * represent *p* ≤ 0.05, ** represent *p* ≤ 0.01, *** represents *p* ≤ 0.001. Data shown are representative of at least three independent biological samples (n ≥ 3).

However, inhibition of pro-survival pathways, ERK or PI3K, under hypoxia condition resulted in induction of *VTN* mRNA expression level, though not significant (**Fig. 5D**). Similarly, Western blotting too demonstrated induction in VTN protein expression levels with both ERK pathway (*p* < 0.01) and PI3K pathway inhibitors (*p* = 0.08) under hypoxia condition, when treated individually, as compared to untreated hypoxic MSCs (**Fig. 5E**). This indicated that VTN expression was negatively regulated via ERK and PI3K pathways under hypoxia condition, which corroborated with our 40°C stress condition data. As a next step, we assessed the viability of WJ-MSCs under hypoxia condition with or without the inhibition of ERK or PI3K pathways, individually. Apoptosis analysis by flow cytometry demonstrated no significant apoptotic change with inhibition of either of the pro-survival pathways under hypoxia condition, with no significant reduction in the viable population percentage (**Fig. 5F, G)**. This again indicated that the upregulated VTN expression might have mediated the survival of hypoxic WJ-MSCs even when the pro-survival pathways were inhibited.

Finally, to validate and affirm the above, we evaluated the effect of ERK pathway inhibition on VTN knocked down WJ-MSCs under hypoxia condition.. VTN knockdown by esiRNA led to reduction in *VTN* mRNA expression (*p* < 0.01), as expected (**Fig. 5H**). Interestingly, corresponding to the reduction in *VTN* expression, apoptosis assay demonstrated a strong reduction in the percentage of viable population from 85.96 ± 1.63 to 66.13 ± 1.69 (*p* < 0.001), with a significant increase in the percentage of late apoptotic population from 4.06 ± 0.49 to 22.53 ± 1.48 (*p* < 0.001) in hypoxic WJ-MSCs treated with ERK pathway inhibitor, as compared to NC siRNA transfected WJ-MSCs under the same treatment (**Fig. 5I, J**).

## Discussion

MSCs have made a significant contribution towards the advancement of cell-based therapy, yet much remains to be explored about their functionality post transplantation. The success of MSC based therapy mainly relies on viability and efficient homing of transplanted MSCs at the inflammation site. However, MSCs get challenged by micro-environmental stress conditions at the inflammatory site, and as a result they undergo massive cell death within a few days of transplantation (21). Thus, understanding the molecular interaction of MSCs with its inflammatory microenvironment would hugely benefit the field of MSC-based regenerative therapy towards the treatment of inflammatory diseases.

Fever is one of the hallmark responses of inflammation and infection, and not much has been investigated on MSC viability and its underlying survival mechanism in response to physiological fever range temperature. As attachment to the ECM is largely known for transmitting survival cues to the cells, exploring the involvement of ECM molecules in the survival of MSCs under inflammatory temperature stress assumes importance. Also, since identifying an appropriate source of MSCs is critical for improved therapeutic efficacy, we utilized human umbilical cord-derived WJ-MSCs, known for better proliferative capacity and immunomodulatory response as compared to adult sources, for our study.

WJ-MSCs, when subjected to temperature stress of 40°C for 48 h, showed an increase in the expression of multifunctional glycoprotein VTN both at mRNA and protein levels. Further, immunofluorescence study depicted widespread increase in relative expression, and change in the subcellular distribution of VTN protein in 40°C exposed WJ-MSCs. In addition, we had also noted that VTN expression increased both in temporal and cell density dependent manner at 40°C, where increase in duration of 40°C stress exposure keeping the cell seeding density constant or increasing cell seeding density keeping the 40°C exposure time same, further induced VTN protein expression. These observations led us to speculate that cellular confluency, or cellular confluency leading to cell growth arrest state could be influencing VTN expression in WJ-MSCs. Extending support to our observation, earlier studies had illustrated cell density-dependent modulation of certain genes like fibronectins, tumor necrosis factor α (TNFα), and insulin-like growth factor I in different cell types. Moreover, the density-dependent increase in the expression of these genes, were shown to be independent of secreted soluble factors (22–24). In contrast, our medium washout experiment indicated that *VTN* expression was regulated via secreted VTN or certain other factors by autocrine or paracrine mode of action in a positive feedback loop.

WJ-MSCs exposed to 40°C upto 48 h did not show any significant apoptotic change, however, they underwent G0/G1 cell cycle arrest. Interestingly, inhibition of prominent prosurvival pathways, ERK and PI3K, at 40°C also did not result in any significant apoptosis, and viability was maintained. This could be attributed to a further upregulation in VTN expression. In addition, with inhibition of ERK or PI3K pathway under 40°C, WJ-MSCs continued to remain in G0/G1 cell cycle arrested state, although there was a reduction in the p53 protein expression (**Supplementary fig. 1C**). These observations overall highlighted the involvement of VTN in maintaining the viability of WJ-MSCs under elevated temperature stress even in the absence of pro-survival pathways, ERK or PI3K, by entering G0/G1 cell cycle arrest.

To reaffirm these observations, VTN knocked down WJ-MSCs were exposed to 40°C temperature stress, following which cell cycle and viability were evaluated. As expected, *VTN* esiRNA transfected WJ-MSCs at 40°C showed a decrease in viable population with increased apoptotic population, moreover, this was also accompanied by reversal in G0/G1 arrest. These data corroborated with our findings that VTN provided pro-survival support to WJ-MSCs via mediating G0/G1 cell cycle arrest. In support, a previous report had shown that growth arrested quiescent human endometrial mesenchymal stem cells were more resistant to heat stress compared to proliferating cells (25). Additionally, earlier report from our lab had also demonstrated the pro-survival effect of VTN in serum deprived WJ-MSCs via mediating G0/G1 cell cycle arrest (18).

Next, treating *VTN* esiRNA transfected WJ-MSCs with inhibitors for the pro-survival pathways under 40°C temperature stress triggered even stronger apoptosis with a prominent reduction in viable population of WJ-MSCs. Furthermore, this reduction in viability was accompanied by lifting of the G0/G1 phase cell cycle arrest. Collectively, these results re-emphasized on the significance of VTN in promoting survival of WJ-MSCs under elevated temperature stress.

In contrast to 40°C temperature stress, on exposure to more favorable physiological condition of hypoxia (2% O_2_), WJ-MSCs displayed not much change in cell morphology or proliferation as compared to control WJ-MSCs. Again, corresponding to these observations, VTN expression remained similar to that of control WJ-MSCs. However, inhibition of the pro-survival pathways, ERK or PI3K, led to strong induction in VTN expression in hypoxia treated WJ-MSCs with no significant change in viability. It is noteworthy that knocking down VTN in hypoxic WJ-MSCs, followed by treatment with ERK pathway inhibitor resulted in reduction in percentage of viable population with corresponding increase in the late apoptotic population. Overall, our study under hypoxia condition further validated the fact that VTN helped to protect and maintain viability of WJ-MSCs exposed to adverse microenvironmental stress conditions. Our observations corroborate with previous findings on the essential role of ECM in cell survival. An earlier report had demonstrated that enrichment of ECM protein like VTN in microenvironment of glioblastoma cells provided cell adhesion mediated survival advantage against chemotherapy (26). Similarly, another study had suggested that using different combinations of ECMs promote better cell survival of human induced pluripotent stem cell-derived endothelial cell under conditions of hypoxia and nutrient deprivation (27).

Finally, to explore the possible involvement of autophagy in VTN’s prosurvival role at 40°C, we studied the expression of p62. p62 is considered as a selective marker to monitor autophagy flux since it binds to LC3 and selectively gets degraded by autophagy (26). Interestingly, p62 protein expression was found to be significantly downregulated at 40°C as compared to control condition WJ-MSCs (**Supplementary fig. 2A-C**). In addition, inhibition of PI3K pathway, negative regulator of VTN expression, led to further downregulation in p62 protein expression, while inhibition of NF-κβ pathway, a positive regulator of VTN expression, exhibited reversal in p62 protein expression (**Supplementary fig. 2A**). VTN knockdown in WJ-MSCs at 40°C also exhibited reversal in p62 protein expression level suggesting VTN’s prosurvival role at 40°C stress being played via autophagy (**Supplementary fig. 2B**, *p* < 0.001). Furthermore, inhibition of autophagy with an autophagy pathway inhibitor, chloroquine, led to upregulation in p62 expression (**Supplementary fig. 2C**, *p* < 0.001) with increase in cell death (**Supplementary fig. 2D**) re-confirming the possible involvement of autophagy. Lending support to our finding, a previous report had also demonstrated that NF-κβ induced autophagy was essential for HeLa cell survival following heat shock treatment (27).

The findings of the present study have adequately established the importance of ECM glycoprotein, VTN, in the survival of WJ-MSCs under the inflammatory temperature stress condition. Furthermore, underlying molecular signaling pathways regulating the expression of VTN under temperature stress condition was elucidated. This knowledge would help in forwarding the understanding regarding MSC survival post transplantation for the treatment of various inflammatory diseases. Further, research on the inflammatory disease-based animal model is warranted to confirm the findings of the study in the context of the role of VTN in the survival of WJ-MSCs.

## Acknowledgments

We thank SERB, DST India and IISER, Kolkata for funding of this work, and CSIR, India for the fellowship of Mr. Umesh Goyal. We are grateful to Dr. Jayanta Chatterjee, Astha, Kalyani, West Bengal, India, for generously providing the umbilical cord samples. We thank Ritabrata Ghosh and Tamal Ghosh for technical assistance with microscopy and flow cytometry, respectively. We are also thankful to Ankita Sen and Srishti Dutta Gupta for their assistance in umbilical cord collection, processing, and cell culture.

## AUTHOR’S CONTRIBUTION

Umesh Goyal: Investigation, analysis of data, validation, writing – Original draft.

Ashiq Khader C.: Investigation and analysis of data.

Malancha Ta: Conceptualisation, supervision, writing-review and editing, funding management.

## References

1) Wang LT, Ting CH, Yen ML, Liu KJ, Sytwu HK, Wu KK, et al. Human mesenchymal stem cells (MSCs) for treatment towards immune-and inflammation-mediated diseases: review of current clinical trials. J Biomed Sci. 2016;23(1):76.

2) Dabrowska S, Andrzejewska A, Janowski M, Lukomska B. Immunomodulatory and Regenerative Effects of Mesenchymal Stem Cells and Extracellular Vesicles: Therapeutic Outlook for Inflammatory and Degenerative Diseases. Front Immunol. 2021;11:591065.

3) Caplan AI. Mesenchymal Stem Cells: Time to Change the Name! Stem Cells Transl Med. 2017;6(6):1445–1451

4) Via AG, Frizziero A, Oliva F. Biological properties of mesenchymal Stem Cells from different sources. Muscles Ligaments Tendons J. 2012;2(3):154–162.

5) Fong CY, Chak LL, Biswas A, Tan JH, Gauthaman K, Chan WK, et al. Human Wharton’s jelly stem cells have unique transcriptome profiles compared to human embryonic stem cells and other mesenchymal stem cells. Stem Cell Rev Rep. 2011;7(1):1–16.

6) Can A, Karahuseyinoglu S. Concise review: human umbilical cord stroma with regard to the source of fetus-derived stem cells. Stem Cells. 2007;25(11):2886–2895.

7) Chen L, Deng H, Cui H, Fang J, Zuo Z, Deng J, et al. Inflammatory responses and inflammation-associated diseases in organs. Oncotarget. 2017;9(6):7204–7218.

8) Evans SS, Repasky EA, Fisher DT. Fever and the thermal regulation of immunity: the immune system feels the heat. Nat Rev Immunol. 2015;15(6):335–349.

9) Goyal U, Ta M. p53-NF-κB Crosstalk in Febrile Temperature-Treated Human Umbilical Cord-Derived Mesenchymal Stem Cells. Stem Cells Dev. 2019;28(1):56–68.

10) Sen A, Ta M. Altered Adhesion and Migration of Human Mesenchymal Stromal Cells under Febrile Temperature Stress Involves NF-κβ Pathway. Sci Rep. 2020;10(1):4473.

11) Leavesley DI, Kashyap AS, Croll T, Sivaramakrishnan M, Shokoohmand A, Hollier BG, et al. Vitronectin--master controller or micromanager? IUBMB Life. 2013;65(10):807–818.

12) Chow S, Di Girolamo N. Vitronectin: a migration and wound healing factor for human corneal epithelial cells. Invest Ophthalmol Vis Sci. 2014;55(10):6590–6600.

13) Scaffidi AK, Moodley YP, Weichselbaum M, Thompson PJ, Knight DA. Regulation of human lung fibroblast phenotype and function by vitronectin and vitronectin integrins. J Cell Sci. 2001;114(Pt 19):3507–3516.

14) Wheaton AK, Velikoff M, Agarwal M, Loo TT, Horowitz JC, Sisson TH, et al. The vitronectin RGD motif regulates TGF-β-induced alveolar epithelial cell apoptosis. Am J Physiol Lung Cell Mol Physiol. 2016;310(11):L1206–1217.

15) Wei F, Tang L, He Y, Wu Y, Shi L, Xiong F, et al. BPIFB1 (LPLUNC1) inhibits radioresistance in nasopharyngeal carcinoma by inhibiting VTN expression. Cell Death Dis. 2018;9(4):432.

16) Hazawa M, Yasuda T, Noshiro K, Saotome-Nakamura A, Fukuzaki T, Michikawa Y, et al. Vitronectin improves cell survival after radiation injury in human umbilical vein endothelial cells. FEBS Open Bio. 2012;2:334–338

17) Bae HB, Zmijewski JW, Deshane JS, Zhi D, Thompson LC, Peterson CB, et al. Vitronectin inhibits neutrophil apoptosis through activation of integrin-associated signaling pathways. Am J Respir Cell Mol Biol. 2012;46(6):790–796.

18) Goyal U, Ta M. A novel role of vitronectin in promoting survival of mesenchymal stem cells under serum deprivation stress. Stem Cell Res Ther. 2020;11(1):181.

19) Goyal U, Sen A, Ta M. Isolation and Molecular Characterization of Progenitor Cells from Human Umbilical Cord. Methods Mol Biol. 2019;2029:1–13.

20) Samal JRK, Rangasami VK, Samanta S, Varghese OP, Oommen OP. Discrepancies on the Role of Oxygen Gradient and Culture Condition on Mesenchymal Stem Cell Fate. Adv Healthc Mater. 2021;10(6):e2002058.

21) Noronha NC, Mizukami A, Caliári-Oliveira C, Cominal JG, Rocha JLM, Covas DT, et al. Priming approaches to improve the efficacy of mesenchymal stromal cell-based therapies. Stem Cell Res Ther. 2019;10(1):131.

22) Perkinson RA, Kuo BA, Norton PA. Modulation of transcription of the rat fibronectin gene by cell density. J Cell Biochem. 1996;63(1):74–85.

23) Fukushima S, Kaneko N, Koiwai O, Koike K. Cell density-dependent regulation of tumor necrosis factor alpha gene expression in a human hepatoma cell line. Int J Oncol. 2007;31(6):1485–1490.

24) Wang L, Adamo ML. Cell density influences insulin-like growth factor I gene expression in a cell type-specific manner. Endocrinology. 2000;141(7):2481–2489.

25) Alekseenko LL, Shilina MA, Lyublinskaya OG, Kornienko JS, Anatskaya OV, Vinogradov AE, et al. Quiescent Human Mesenchymal Stem Cells Are More Resistant to Heat Stress than Cycling Cells. Stem Cells Int. 2018;2018:3753547.

26) Yu Q, Xiao W, Sun S, Sohrabi A, Liang J, Seidlits SK. Extracellular Matrix Proteins Confer Cell Adhesion-Mediated Drug Resistance Through Integrin α v in Glioblastoma Cells. Front Cell Dev Biol. 2021;9:616580.

27) Hou L, Coller J, Natu V, Hastie TJ, Huang NF. Combinatorial extracellular matrix microenvironments promote survival and phenotype of human induced pluripotent stem cell-derived endothelial cells in hypoxia. Acta Biomater. 2016;44:188–199.

28) Yoshii SR, Mizushima N. Monitoring and Measuring Autophagy. Int J Mol Sci. 2017;18(9):1865.

29) Nivon M, Richet E, Codogno P, Arrigo AP, Kretz-Remy C. Autophagy activation by NFkappaB is essential for cell survival after heat shock. Autophagy. 2009;5(6):766–783.

