## Supplemental Figure for "Vitronectin mediates survival of human WJ-MSCs under inflammatory temperature stress via cell cycle arrest"

**Supplementary fig 1:**

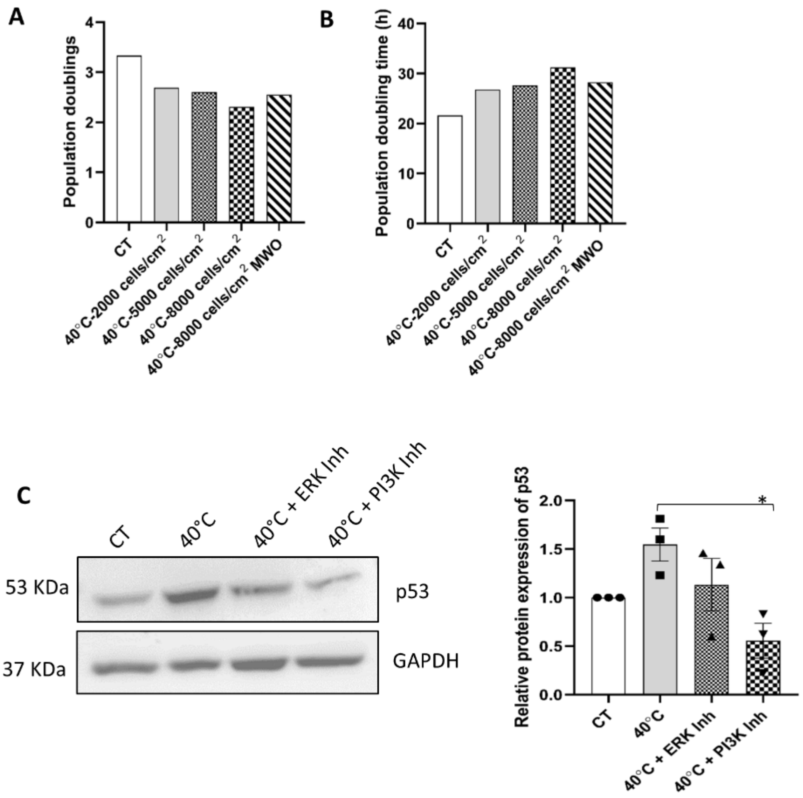


**Supplementary figure 1:** Effect of medium washout (MWO) experiment on population doubling time and population doublings. WJ-MSCs plated at high seeding density of 8000 cells/cm$^2$ under 40°C, when subjected to medium change every 12 h showed a rescue, that is (**A**) an increase in the number of population doublings and (**B**) decrease in population doubling time as compared to the MSC sample seeded at 8000 cells/cm$^2$ but without any medium change (n = 2).

Effect of ERK or PI3K pathway inhibition on p53 protein expression in WJ-MSCs exposed to 40°C. (**C**) p53 protein expression was detected in WJ-MSCs exposed to 40°C for 48 h in the absence or presence of ERK pathway inhibitor, FR180204 or PI3K pathway inhibitor, LY294002. Representative Western blotting images from three independent biological samples are displayed. Band densities were quantified and plotted relative to GAPDH expression used as loading control. Each bar represents mean ± SEM.* represent $p \leq 0.05$.

**Supplementary fig 2:**

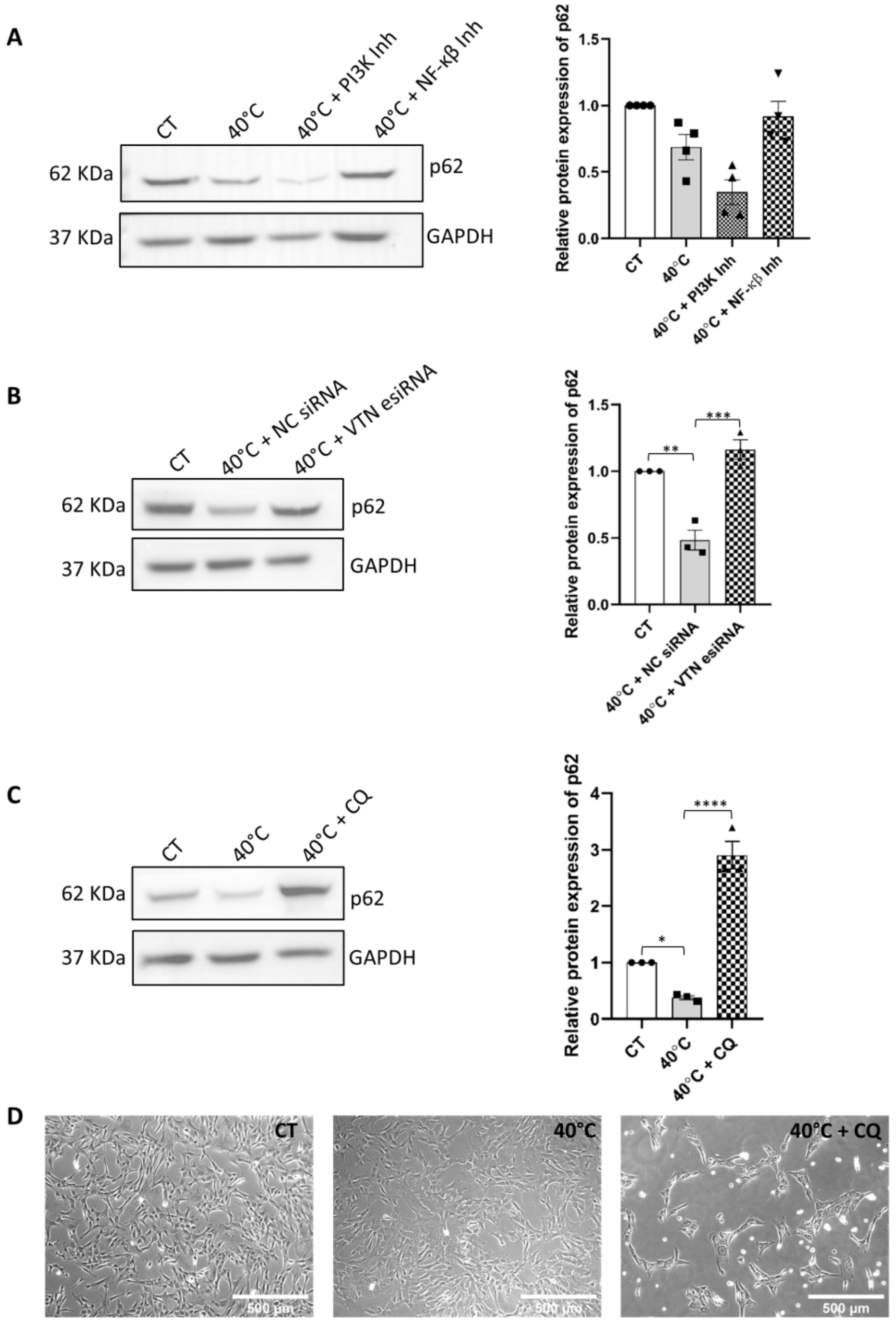


**Supplementary figure 2:** Exploring involvement of autophagy pathway by detecting expression level of p62. (**A**) WJ-MSCs were subjected to 40°C in the absence or presence of small molecule inhibitors for PI3K and NF-κβ pathways, individually, for 48 h. p62 protein expression was detected and representative Western blotting images from four independent biological samples are displayed. Band densities were quantified and plotted relative to

GAPDH expression, used as loading control. (**B**) WJ-MSCs were transfected with *VTN* esiRNA, or NC siRNA and were exposed to 40°C for 48 h. p62 protein expression was detected and representative Western blotting images from three independent biological samples are displayed. Band densities were quantified and plotted relative to GAPDH expression, used as loading control. (**C**) For autophagy inhibition study, WJ-MSCs were treated with chloroquine (CQ) at a concentration of 25 μM, and p62 protein expression was assessed. Representative Western blotting images from three independent biological samples are displayed. Band densities were quantified and plotted relative to GAPDH expression, used as loading control. (**D**) Representative phase-contrast morphology images of control and 40°C treated WJ-MSCs, in the absence or presence of chloroquine, under 10X magnification (n = 3). Each bar represents mean ± SEM. * represent $p \leq 0.05$, ** represent $p \leq 0.01$, *** represents $p \leq 0.001$ and **** represents $p \leq 0.0001$. Data shown are representative of at least three independent biological samples (n ≥ 3).
